# Arabidopsis cobra protein binds to glucan chains with higher affinity than to cellulose

**DOI:** 10.1101/050427

**Authors:** Nadav Sorek, Chris Somerville

## Abstract

Because of several anomalies in a publication that we coauthored, concerning the interaction of the Arabidopsis cobra protein with crystalline cellulose (Avicel) and individual glucan chains (cellohexaose) we reexamined the data and had an independent laboratory redo a key experiment. The results show that COBRA binds to cellohexaose preferably over binding to Avicel, confirming our previous report.

## Introduction

Mutations in the Arabidopsis *COBRA* gene have defects in cell wall biosynthesis but the specific function of the corresponding protein is unknown (Roudier et al., 2005). Expression of the *COBRA* gene is strongly correlated with expression of the *CESA* genes that are thought to be the catalytic components of the cellulose synthase complex (Persson et al., 2005).

In a paper published in 2014 (Sorek et al., 2014) we measured the interaction of COBRA with individual glucan chains (cellohexaose) by Isothermal Calorimetry (ITC). We also performed a competition assay, in which COBRA binding to crystalline cellulose (Avicel) was inhibited by a range of concentrations of cellohexaose. The results were interpreted as indicating that COBRA binds cellohexaose with higher affinity than Avicel. Subsequently, an observer noticed anomalies in the ITC chromatogram and the gel blot. To address concerns about the validity of the published results, we revisited the results of the ITC and asked an independent researcher to repeat the competition assay of COBRA binding to cellohexaose and cellulose.

## Results

### Isothermal titration calorimetry of cobra binding to cellohexaose

In the Sorek et al., (2014) paper there appears to be a discontinuity of the ITC tracing from the background grid. We reanalyzed the raw data and contacted the ITC manufacturer (TA instruments) for technical explanation. Upon reanalyzing the original data, we did not observe any anomalies in the plot (Figure 1). Reanalysis of the raw data yielded the same kinetic parameters. The interpretation of TA Instruments was that the anomaly was due to a failure in the graphic card in the previously published plot.

**Figure 1.**
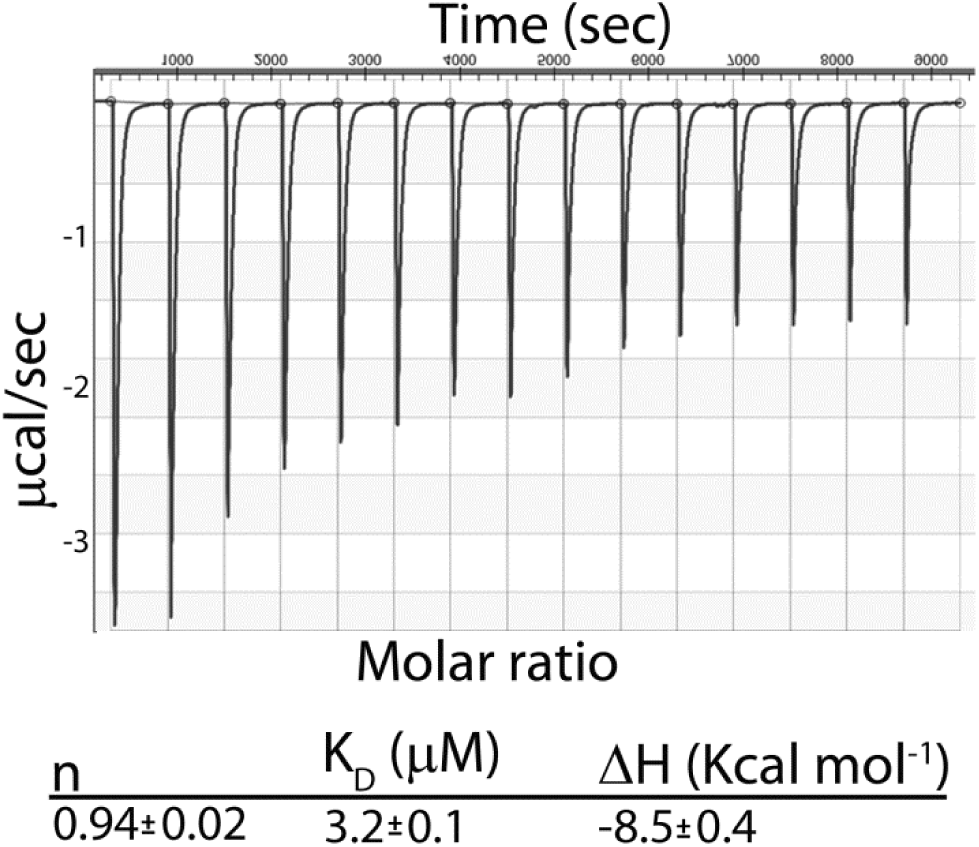
Isothermal Calorimetry chromatogram of COBRA binding to cellohexaose. The upper plot is a screenshot of the raw data from the binding assay between COBRA protein and cellohexaose. For convenience, the derived kinetic parameters are presented.

## COBRA binding to cellohexaose and cellulose

In the Sorek et al., (2014) paper we tested the binding preference of COBRA to cellulose and glucan chains. The image of the gel blot was saved as a JPEG file, which when printed, exhibited anomalies under certain exposure conditions. We further repeated the experiment several times and got the same results, without the anomalies present in the published paper. To remove any doubt, we asked an independent postdoctoral researcher outside the lab (Dr. Wakao, UC Berkeley) to repeat the experiment, and these results are presented in figure 2.

**Figure 2.**
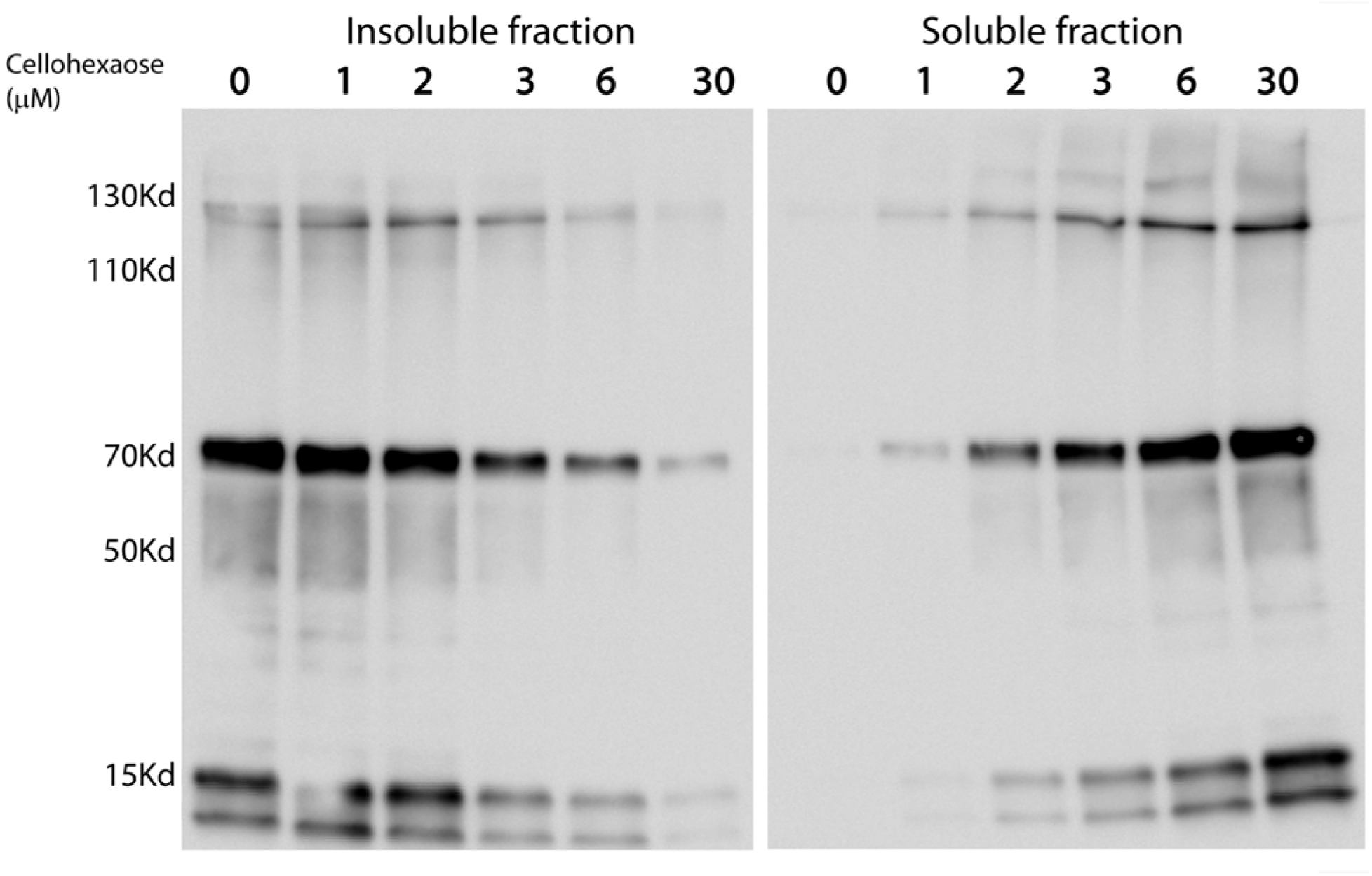
cobra binds to cellohexaose with higher affinity than to cellulose. COBRA protein was incubated with Avicel and increasing concentrations of cellohexaose. Without cellohexaose, COBRA binds to Avicel with negligible detection in the soluble fraction. As the concentrations of the cellohexaose increase, COBRA was detected predominantly in the soluble fraction, suggesting higher affinity of COBRA for glucan chains than for cellulose.

Competition assays were carried out using pulldown assays. COBRA was incubated for 2 h at 21 °C with Avicel in the presence of increasing concentrations of cellohexaose (0 to 30μM). The washed pellet (insoluble fraction) represents COBRA that was bound to Avicel, and the soluble fraction represents COBRA protein that was not bound to Avicel (figure 2). With no cellohexaose present, COBRA is bound to Avicel and no soluble protein can be detected. COBRA KD of binding cellohexaose is 3.2 μM, and with 3 μM cellohexaose, more protein was detected in the soluble fraction compared to the insoluble, Avicel bound fraction. With 6 μM cellohexaose the majority of COBRA was soluble and with 30 μM of cellohexaose, COBRA could be hardly detected in the insoluble fraction.

The results in figure 2 match the results published in Sorek et al., (2014). In both cases, increasing concentrations of cellohexaose are able to surpass the binding of COBRA to cellulose. Therefore, we believe that the hypothesis that COBRA might act at the initial stage of cellulose synthesis where individual glucan chains are assembled into cellulose fiber is supported by the available data.

## Material and methods

### Isothermal Titration Calorimetry

The chromatogram shown in figure 1 is for the same experiment that was published by Sorek et al., (2014). Isothermal titration calorimetry experiments were performed in NanoITC2G (TA Instruments). Measurements were made at 21 °C. The proteins were dialyzed extensively against 50 mm Tris-Cl buffer, pH 7.5, and the cellohexaose was dissolved in the same buffer to minimize heats of dilution. Titrations were carried out in triplicate, and the errors are the mean ± S.D. of these replicates. During a titration experiment the 50 μm protein sample stirred at 250 rpm in a 1.43-ml reaction cell that was injected with 15 successive 15-μl aliquots of 750 μm cellohexaose at 300-s intervals. Integrated heat effects, after correction for heats of dilution, were analyzed by Sigmaploid dose-response + Hill slope (SigmaPlot). The fitted data yielded the dissociation constant (Kd), number of binding sites on the protein (n), and the enthalpy of binding (ΔH).

### COBRA, Avicel, and Cellohexaose Competition Assay

The experiment was repeated with the exact conditions as in Sorek et al., (2014). Proteins were incubated with Avicel PH-101 (Sigma, 11365) (1 mg/ml) in 20 mm Tris-Cl buffer, pH 8, and 150 mM NaCl for 120 min. Avicel recovered by centrifugation was washed 3 times with 20 mm Tris-Cl buffer, pH 8, and 150 Mm NaCl. Avicel-bound proteins were released by heating in SDS sample buffer for 5 min at 100 °C and subjected to SDS-PAGE.

## Acknowledgements

We would like to thank Dr. Setsuko Wakao for performing the binding assay.

